# Probabilistic model discovery reveals distinct constitutive behavior of kidney cortex and medulla

**DOI:** 10.64898/2026.09.12.751158

**Authors:** Vilde Ellingsen, Skyler R. St. Pierre, Ellen Kuhl

## Abstract

The kidney is a critical soft-tissue organ responsible for blood filtration. Accurate constitutive models of the kidney are essential to predict tissue deformation and stress, yet existing models prescribe the strain energy function a priori, largely treat tissue variability as deterministic, and do not distinguish between cortex and medulla. Here we use Gaussian constitutive artificial neural networks to discover probabilistic strain energy functions for kidney cortex and medulla from tension, compression, and shear experiments. The medulla is approximately twice as stiff as the cortex across all three modes: The effective Young’s moduli are 3.41, 5.04, and 4.48 kPa for the medulla and 1.43, 2.32, and 2.87 kPa for the cortex in tension, compression, and shear. Both regions display pronounced tension–compression asymmetry. When trained on all three modes, the network discovers strain energy functions in the second invariant alone over the tested deformation range. The cortex model selects two exponential *I*_2_ terms, while the medulla model selects an additional linear *I*_2_ term. Gaussian external weights propagate specimen-to-specimen variability into closed-form probabilistic stress predictions. Our region-specific probabilistic models capture the mechanical heterogeneity and variability of the kidney and enable more realistic simulations of kidney deformation and stress for surgical planning, needle interventions, and renal trauma. All data, source code, and examples are available at https://github.com/LivingMatterLab/CANN.

## 1. Motivation

Kidney tissue is extremely soft and porous which makes it vulnerable to traumatic injury [11, 18]. Disease and infection can alter the mechanical properties of the kidney together with its function. Naturally, linking kidney biomechanics and function holds potential for non-invasive diagnostics and early disease detection [15]. Despite their importance, the mechanics of the kidney remain inconsistently characterized, and the reported properties vary largely between studies [19, 35]. Differences in experimental protocol, modeling assumptions, and the preselected constitutive model contribute to this spread [4, 10]. Three gaps follow: First, every constitutive study of the kidney pre-selects the functional form of the model which imposes a user bias. Second, existing kidney models report either a single parameter set or a mean and standard deviation of parameters, and neither propagates the specimen-to-specimen scatter into the predicted stress. A model that also predicts the variance gives a more complete picture of the tissue [4, 31]. Third, the cortex and the medulla differ in structure and function, yet no constitutive model distinguishes the two regions.

The kidney has distinct regions, each with its own structure and function. Together the cortex and the medulla form the parenchyma, the functional tissue that holds the nephrons (Fig. 1). In both regions, interstitial cells embedded in an extracellular matrix fill the space between the tubules and blood vessels [25]. The *cortex*, the outer layer, begins blood filtration and performs most reabsorption. Densely packed glomeruli– capillary bundles inside cup-shaped Bowman’s capsules–and tangled tubules give it a granular, largely isotropic microstructure. The *medulla*, the inner region, forms pyramids in which radially oriented loops of Henle, collecting ducts, and vasa recta impose a directional, anisotropic microstructure [1]. Here the kidney concentrates urine and conserves water [1].

**Figure 1:**
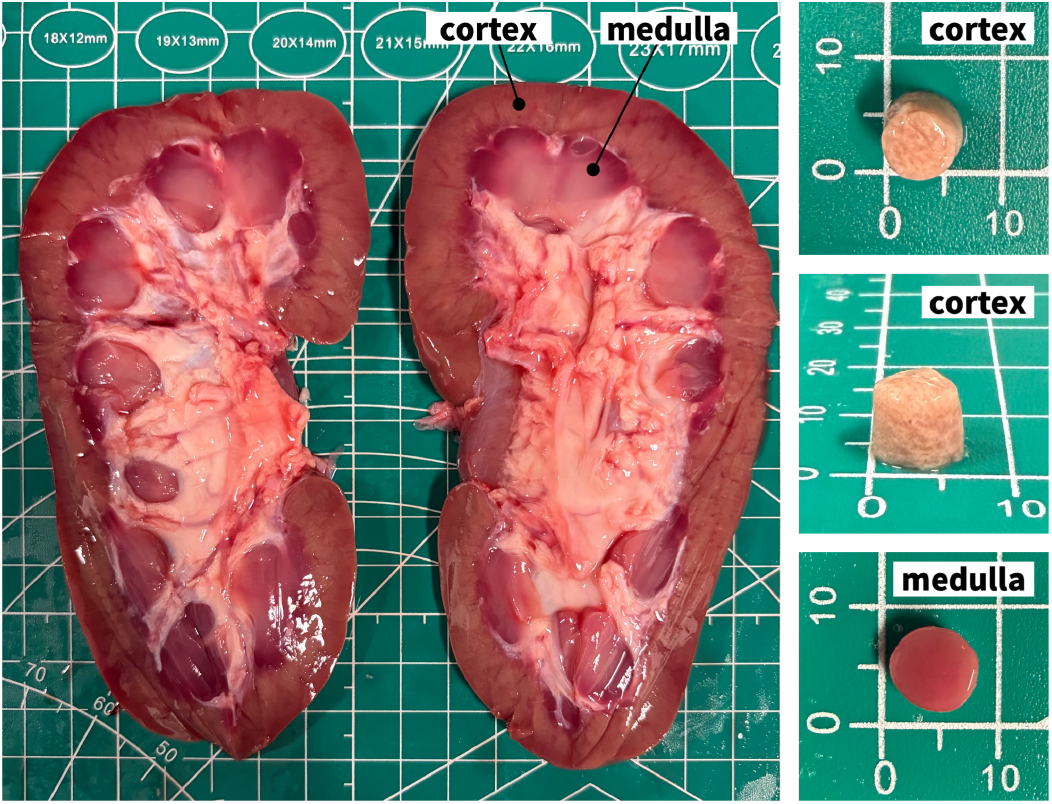
Kidney cross-section and test specimens. Bisected porcine kidney with the outer cortex and inner medulla (left). Cylindrical punch specimens cored from each region, sized against a millimeter grid (right).

Most regional stiffness data for the kidney originate from imaging rather than from mechanical testing. Two methods dominate: ultrasound shear-wave elastography and acoustic radiation force impulse imaging [14, 29, 47, 48, 53], and magnetic resonance elastography or tomoelastography [8, 15, 30, 33, 50, 51]. Both characterize linear shear viscoelasticity rather than the finite stress–stretch response. While a recent study tests both cortex and medulla in compression and shear, it neither compares nor models the two regions explicitly [42]. Among research that does compare both regions within the same study, the ordering is unclear: some report a stiffer cortex [29, 48, 53], a stiffer medulla [8, 15, 47], no difference [50, 51], and other non-monotonic orderings depending on direction, region, and probe orientation [14, 30, 33]. These studies also span different species and preparations, and measurement conditions account for additional spread [8, 14, 33]. In nearly every study, regional comparisons are a byproduct of a different objective, either the discrimination of healthy from diseased tissue or the development of an imaging method. No generally accepted baseline exists, and no study compares the two regions under quasi-static, finite deformations.

Several groups have modeled the constitutive behavior of kidney tissue. In every existing study, the authors first preselect a constitutive model and then fit its parameters to experimental data. Popular forms include the Blatz-Ko model [11], Mooney–Rivlin for the elastic response [36], Ogden for the capsule with Mooney–Rivlin for the cortex [49], linear elasticity [20], reduced-polynomial viscoelasticity [37], a phenomenological tanh stress–strain relation [19], linear viscoelasticity [34], fractional viscoelasticity [35], and damage-dependent viscoelasticity [52]. Across the existing literature, the choice of the functional form precedes the data and injects the modeler’s bias into the result. Automated model discovery reverses that order: a custom-designed, physically-grounded network discovers the model and parameters directly from data [26].

Constitutive models can improve finite element simulations of renal surgery, trauma, and disease. These simulations inherit the uncertainty of their material model. Upper and lower bounds on a simulated outcome therefore require an uncertainty-aware constitutive model [41]. Studies that report linear or viscoelastic moduli sometimes report them as a mean with ± standard deviation across specimens [19, 20, 34, 35], whereas hyperelastic fits often only report a single deterministic parameter set [11, 36, 37, 49, 52]. In neither case does the scatter enter the model. Given the large inter-specimen variability of biological tissue, a constitutive model should predict the variance alongside the mean [28, 31]. This can also provide a healthy-tissue baseline for disease quantification, since the mechanics of diseased kidney tissue can differ substantially from healthy tissue [30, 48, 53].

Every machine-learning strategy for constitutive modeling is a tradeoff between flexibility, interpretability, physical consistency, data requirements, and uncertainty quantification [13]. Feedforward and physics-informed networks fit stress–strain data flexibly, but lack interpretability, and a loss penalty enforces polyconvexity only weakly [13]. Input-convex networks and neural ordinary differential equations guarantee convexity by construction, yet remain black boxes [22, 45, 46]. Model-free and sparse-regression methods such as EUCLID need dense or full-field data [12, 21], and Gaussian process regression returns a posterior variance but no closed-form equation without a further distillation step [3]. Constitutive artificial neural networks discover an interpretable, polyconvex strain energy function from sparse homogeneous tests [26, 46]. Their Bayesian extension quantifies uncertainty through variational weight posteriors, but requires pre-selecting a prior [28]. Gaussian constitutive artificial neural networks (GCANN) need no prior, and map the weight mean, variance, and correlation onto a closed-form stress distribution [31]. We therefore adopt Gaussian constitutive artificial neural networks for kidney tissue.

Here we address all three gaps. We perform tension, compression, and simple shear tests on porcine kidney cortex and medulla, and train Gaussian constitutive artificial neural networks on these data to discover both strain energy functions and predictive uncertainty for each region. Section 2 describes the experiments and the neural networks, Section 3 reports the discovered models and their fit across the different loading modes, and Section 4 interprets the results and their limitations.

## 2. Methods

### 2.1. Experimental Methods

We prepare samples from three porcine kidneys purchased intact from a commercial grocery store. Immediately after purchase, we freeze the kidneys at − 20 °C until testing. We then thaw them in a refrigerator for 12 h at 4 °C before testing. We extract cylindrical samples with an 8 mm surgical biopsy punch. We trim the ends flat with a surgical scalpel to a height of 6–9 mm. By visual inspection, we assign each sample to the cortex or medulla, and do not control for sample orientation within the kidney. After extraction, we immerse each specimen in phosphate-buffered saline and keep it immersed until testing. We weigh each specimen before and after immersion and report the weight change (Fig. 6). We measure the diameter of each sample immediately before testing and use it to compute the stress.

We test 20 samples, 10 per region, on a TA Instruments Discovery HR-20 rheometer (New Castle, DE, USA) fitted with 180-grit sandpaper grips. Our protocol adapts established testing protocols for ultrasoft biological tissues [6, 9]. To improve glue adhesion, we lightly dry the top and bottom faces of the immersed sample. Then we apply a small amount of cyanoacrylate adhesive (Loctite Super Glue Ultra Gel Control, Henkel Consumer Adhesives) to the upper face, and press the sample against the upper plate for 30 s until the glue sets (Fig. 2, left). We then lower the sample onto the lower plate over a second thin layer of adhesive and confirm contact visually (Fig. 2, middle). The glue then cures for 60 s, and while we configure the test (Fig. 2, right).

**Figure 2:**
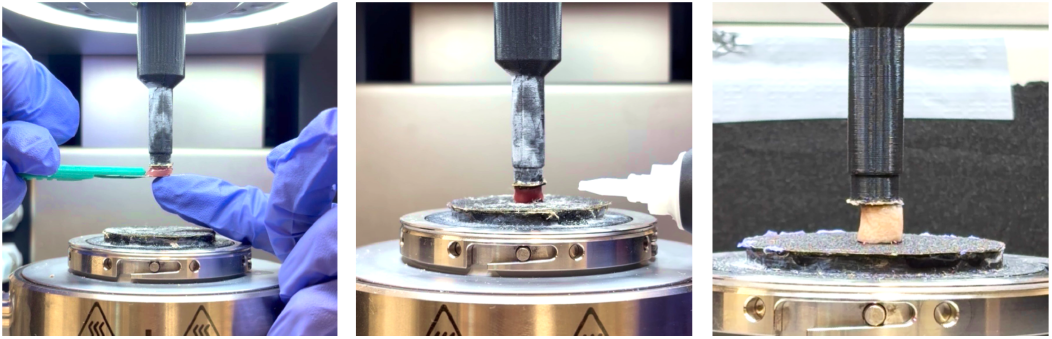
Specimen mounting. Cyanoacrylate glue adheres the specimen to the upper geometry (left) and to the lower plate (middle). To prevent dehydration, we mist the mounted specimen (right) with phosphate-buffered saline.

Although it is well accepted that dehydration alters the mechanical properties of the sample and makes it more rigid, there is no standardized guidance for sample hydration. To mimic the fully hydrated in vivo state, some researchers recommend a bath of saline solution during testing [38], but wetted specimens typically recover their original hydrated properties upon re-hydration [7]. With our current setup, we cannot fully sub-merge the sample during testing. Instead, we keep each sample submerged in phosphate-buffered saline from extraction until testing, and mist it before testing and during the inter-mode holds. We visually confirm that the sample never dries.

We zero all forces after hydration and before loading, and conduct all tests at ambient room temperature. We test each sample in a fixed sequence of compression, tension, and simple shear (Table 1). All tests use a quasi-static load rate, axial 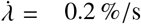 and shear 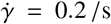. Between modes, we hold the sample unloaded for 60 s. We record the sample height for each test to compute the stretch *λ* and the amount of shear *γ*. We treat the first cycle of each mode as preconditioning and use only the last complete loading and unloading cycle for training. This procedure gives a repeatable response across the three modes tested consecutively on the same sample.

**Table 1:** Loading protocol for compression, tension, and shear testing. Summary of the loading modes, cycle counts, and target deformations applied to each sample. In compression and tension, we ramp the stretch linearly to the target value, then reverse the ramp linearly back to the undeformed configuration, and we repeat this cycle three times. In shear, we ramp the shear strain linearly to *γ* = 0.15 and back to zero twice, then apply a single ramp to *γ* = 0.30. We hold the sample unloaded for 60 s between consecutive modes to let the tissue recover before the next mode begins.

| step | mode | cycles | load / unload path |
| --- | --- | --- | --- |
| 1 | compression | 3× | load $\lambda$ : 1 → 0.9, unload $\lambda$ : 0.9 → 1 |
| 2 | relax | — | no load, 60 s |
| 3 | tension | 3× | load $\lambda$ : 1 → 1.1, unload $\lambda$ : 1.1 → 1 |
| 4 | relax | — | no load, 60 s |
| 5 | shear | 2× | load $\gamma$ : 0 → 0.15, unload $\gamma$ : 0.15 → 0 |
| 6 | shear | 1× | load $\gamma$ : 0 → 0.30 |

### 2.2. Stress and Strain Analysis

We process the rheometer output to obtain the stretch *λ* and the first Piola–Kirchhoff stress *P* for the compression and tension modes, and the amount of shear *γ* and the shear stress *τ* for the shear mode,

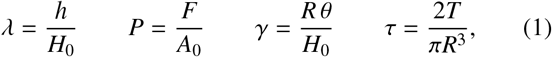

where *H*_0_, *R*, and *A*_0_ = *πR*^2^ denote the undeformed sample height, radius, and cross-sectional area. The deformed quantities *h, F, T*, and *θ* denote the measured height, axial force, torque, and angle of rotation. We define the elastic response as the average of the loading and unloading branches of the third cycle in compression and tension and the second cycle in shear. Before averaging, we interpolate both branches of the selected cycle onto a common deformation grid. For each mode and both regions, we resample each averaged curve to 200 uniformly spaced points for model discovery and to 17 discrete points for the stretch-stress plots. Within the linear regime, we determine the tensile and compressive Young’s moduli *E*_ten_ and *E*_com_ from linear regressions of stress against engineering strain, and the shear Young’s modulus *E*_shr_ = 2(1 + *v*) *G*_shr_ = 3*G*_shr_ from linear regression of shear stress against shear strain, assuming incompressibility, *v* = 0.5.

### 2.3. Continuum Mechanics

The motion ***x*** = φ(***X***) maps each material point ***X*** in the reference configuration to its position ***x*** in the deformed configuration. We describe the local deformation with the deformation gradient

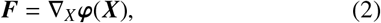

and multiplying ***F*** by its transpose ***F***^t^ gives the symmetric right Cauchy–Green deformation tensor ***C***,

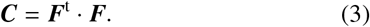

To characterize the deformation of an isotropic material, we introduce the first and second invariants,

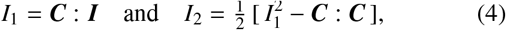

where ***I*** is the identity tensor. We assume the tissue is perfectly incompressible, *J* = det(***F***) = 1, so *I*_1_ and *I*_2_ alone characterize the deformation state. The Piola stress ***P*** relates the force in the current configuration to the undeformed reference area. For a hyperelastic material, ***P*** derives from the strain energy function *ψ*,

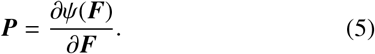

The Piola stress ***P*** then follows from eq. (5) by the chain rule across the invariants [17],

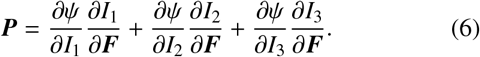

To enforce incompressibility, we replace the third-invariant term with a pressure term, *∂ψ/∂I*_3_ ·*∂I*_3_*/∂****F*** = − *p* ***F***^−t^, where the Lagrange multiplier *p* maintains *J* = 1. The Piola stress becomes

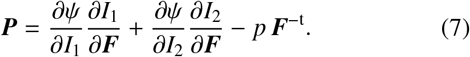

For uniaxial tension and compression, the deformation gradient is ***F*** = diag(*λ, λ*^−1*/*2^, *λ*^−1*/*2^), where incompressibility requires *λ*_2_ = *λ*_3_ = *λ*^−1*/*2^. The invariants reduce to *I*_1_ = *λ*^2^ + 2*λ*^−1^ and *I*_2_ = 2*λ* + *λ*^−2^, and the axial first Piola stress is

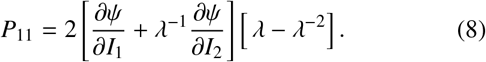

For simple shear with engineering shear strain *γ*, the deformation gradient is ***F*** = ***I*** + *γ* ***e***_1_ ⊗ ***e***_2_. Thus, the invariants equal *I*_1_ = *I*_2_ = 3 + *γ*^2^, and the shear stress is

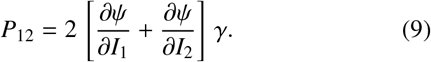

Since *I*_1_ = *I*_2_ throughout simple shear, we can only identify the sum *∂ψ/∂I*_1_ + *∂ψ/∂I*_2_ from shear data alone. Tension or compression data must enter the fit simultaneously to separate the *I*_1_ and *I*_2_ contributions.

### 2.4. Constitutive Neural Networks

Constitutive artificial neural networks discover constitutive models that satisfy thermodynamic consistency, polyconvexity, material objectivity, and material symmetry by design [26]. The network discovers a strain energy function *ψ* in terms of the invariants of the right Cauchy–Green tensor ***C***, and the discovered invariants encode information about the material. We treat kidney tissue as incompressible, such that the *I*_3_ = det ***C*** = 1 is constant, and as isotropic, such that *ψ* depends only on the first and second invariants *I*_1_ and *I*_2_.

The first network architecture, which we call the two-activation architecture, uses two convex activation functions, the identity (°) and the exponential (exp(°)−1) (Fig. 3). The network has *n* = 8 terms and 16 network weights, eight internal 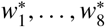 and eight external *w*_1_, …, *w*_8_, and its strain energy function is

**Figure 3:**
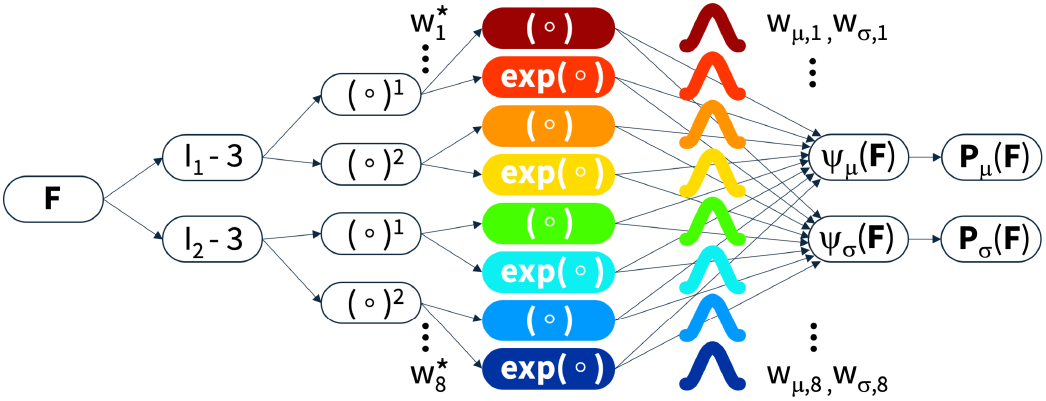
Gaussian constitutive artificial neural network. The neural network maps the first and second invariants *I*_1_ and *I*_2_ onto their first and second powers (°)^1^ and (°)^2^ and applies the identity and the exponential function (°) and exp(°) − 1 to obtain the strain energy *ψ*. A probabilistic output layer returns a Gaussian distribution over the predicted stress ***P***.

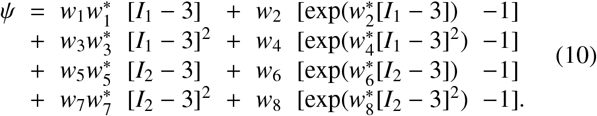

With non-negative weights, 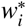, *w*_*i*_ ≥ 0, the two convex, monotonically increasing activation functions guarantee a polyconvex *ψ*.

The second architecture, which we call the enhanced logarithmic architecture, adds a third activation function (ln(1 + °)) and therefore has *n* = 12 terms and 24 network weights, 12 internal 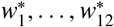, 12 external *w*_1_, …, *w*_12_ (Fig. 4). Its strain energy function is

**Figure 4:**
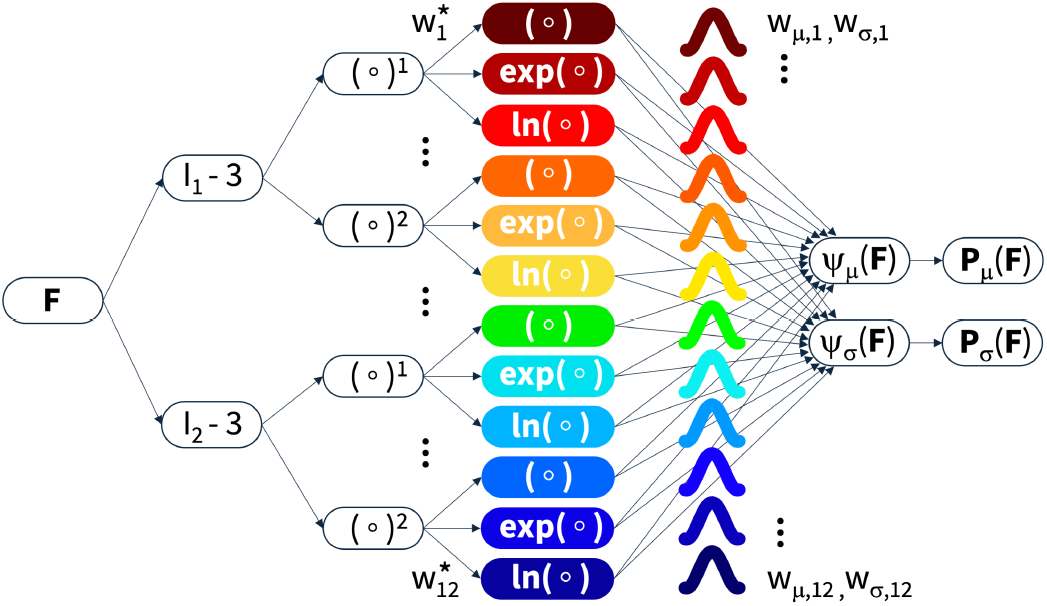
Enhanced Gaussian constitutive artificial neural network. The enhanced neural network maps the first and second invariants *I*_1_ and *I*_2_ onto their first and second powers (°)^1^ and (°)^2^ and applies the identity, the exponential function, and the natural logarithm (°), exp(°) −1, and ln(1+°) to obtain the strain energy *ψ*. A probabilistic output layer returns a Gaussian distribution over the predicted stress ***P***.

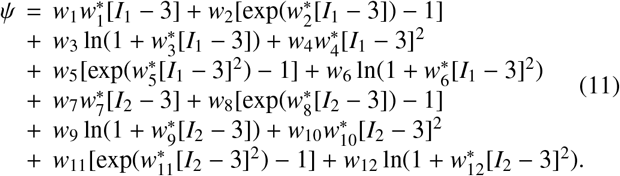

The third activation function (ln(1 + °)) is monotonically increasing, but concave. However, it gives the network an added flexibility that the linear and exponential term cannot provide, so we include it. Equation (11) still guarantees a stress-free reference configuration and objectivity, but no longer polyconvexity. For both networks, each term contributes to the strain energy function independently and no term couples the two invariants. The first Piola–Kirchhoff stress ***P*** follows directly from the discovered strain energy function,

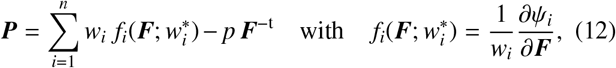

where the Lagrange multiplier *p* enforces incompressibility.

### 2.5. Gaussian constitutive artificial neural networks

A deterministic constitutive neural network returns one stress prediction per deformation state and learns it by minimizing the mean squared error (MSE) between predictions and data. For data that scatter about the prediction with a constant variance, the mean squared error equals the negative log-likelihood (NLL) of the weights up to a constant factor and offset, so the network recovers the mean of the data and carries no information about their spread [31, 40]. Gaussian neural networks [40] predict a variance *σ*^2^ alongside the mean *μ*, both as functions of the input and the weights, and the equivalence between MSE and NLL no longer holds. Each residual now enters the loss weighted by (1*/σ*^2^), a (ln *σ*^2^) term penalizes an inflated variance, and the negative log-likelihood replaces the mean squared error as the training objective.

A Gaussian constitutive artificial neural network predicts both stress mean and stress variance as functions of the deformation gradient ***F***. We adapt the independent Gaussian network model [31] for the isotropic case with uniaxial stretch and simple shear. The network separates deterministic internal weights from probabilistic external weights (Fig. 3). The internal weights carry no variance, and keeping them deterministic preserves the interpretability of the model. The external weights alone introduce uncertainty, and we treat them as independent, so their covariance matrix is diagonal. Correlated weights could further improve predictive performance [31], but we do not consider them here.

The Piola stress follows eq. (12). We restate the derivation of for a single stress component *P* at a given deformation gradient ***F***. In tension and compression, *P* is the axial Piola stress *P*_11_ of eq. (8). In shear, *P* is the shear stress *P*_12_ = *τ* of eq. (9). Both are linear in the external weights,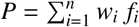. We model the external weights as Gaussian random variables [31],

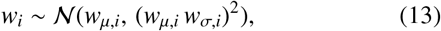

where *w*_*μ,i*_ is the mean weight and *w*_*σ,i*_ scales its standard deviation relative to that mean. We normalize each weight by its mean, 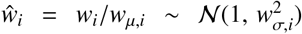, such that ***ŵ*** ∼ *N*([1, …, 1], Σ) with 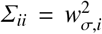, and define the stress as 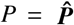· ***ŵ*** with the constant vector 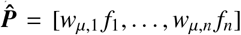. Since 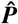 is deterministic at a given ***F*** and ***ŵ*** follows a multivariate normal distribution, *P* is normally distributed. Its mean is

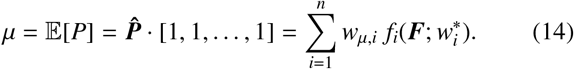

The independent model uses a diagonal covariance matrix, 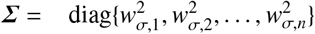, so the variance simplifies to

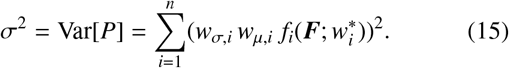

Polyconvexity requires non-negative external weights, *w*_*i*_≥ 0. With non-negative means, *w*_*μ,i*_ ≥ 0, the constant-exponential architecture (Fig. 3) guarantees polyconvexity in the mean. A Gaussian weight always retains a finite probability of turning negative, so we cannot enforce non-negativity exactly. Instead we bound the probability that a weight is negative. We restrict the diagonal of the covariance matrix [31],

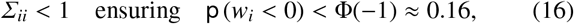

where Φ is the cumulative distribution function of the standard normal.

### 2.6. Training Objective

We estimate the deterministic model parameters 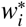, *w*_*μ,i*_, and *w*_*σ,i*_ by maximum likelihood. The external weights *w*_*i*_ remain random variables. The likelihood gives the probability of the measured stresses under a given parameter set. Rather than maximizing the likelihood directly, we minimize the negative log-likelihood. At each data point *k, P*_*k*_ denotes the measured stress, and *μ*_*k*_ and 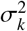 denote the predicted mean and variance, with eqs. (14) and (15) evaluated at ***F***_*k*_. The negative log-likelihood over *N* data points is [31]

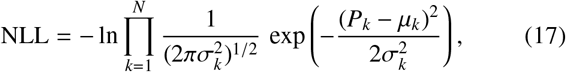

which simplifies to

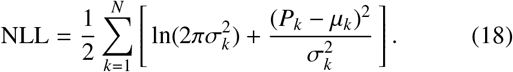

To compare fits across modes and training sets, we report the excess negative log-likelihood, eNLL = NLL − NLL_0_, where NLL_0_ is the value eq. (18) takes when *μ*_*k*_ and 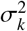 equal the specimen mean and variance of the data at each point. This floor is the lowest negative log-likelihood any Gaussian can reach, so eNLL = 0 is a perfect fit and larger values are worse. To control the number of active parameters and prevent overfitting, we add an *L*_*p*_ regularization term,

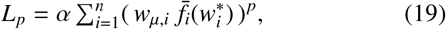

where 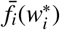 is the stress contribution of term *i* at unit weight integrated over the training deformation range [31]. A value of *p* = 0.5 promotes sparsity [32], so we adopt it. The total loss is *L* = NLL + *L*_*p*_. We vary the penalty parameter *α* from 0 to 1 to balance a low loss against model interpretability [31, 32].

We train the independent, regularized Gaussian constitutive artificial neural networks with the variance constrained such that *Σ*_*ii*_ *<* 1 (eq. (16)). Starting from random parameter initialization, we minimize the negative log-likelihood loss (eq. (18)) with the ADAM optimizer, at a learning rate of 0.001 and a batch size of 1,000. We pre-train for 2,000 epochs without regularization, then apply the *L*_*p*_ penalty (eq. (19)) for the next 3,000 epochs, for 5,000 epochs total [31]. To select *α*, we sweep *α* over the range from zero to one. We choose *α* = 0.01 for both the cortex and medulla, the value that gives the fewest terms without a notable increase in eNLL.

## 3. Results

### 3.1. Experimental response

The medulla is stiffer than the cortex in every mode (Table 2). Linear fits near the reference configuration yield effective moduli of 1.43, 2.32, and 2.87 kPa for the cortex and 3.41, 5.04, and 4.48 kPa for the medulla in tension, compression, and shear. The gap is largest in compression and smallest in shear (Fig. 5). The medulla peak stress reaches 3.97 times the cortex value in compression with − 2.56 vs. − 0.64 kPa at *λ* = 0.9, 2.37 times in tension with 0.37 vs. 0.16 kPa at *λ* = 1.1, and 1.59 times in shear with 0.18 vs. 0.11 kPa at *γ* = 0.11. Both regions show pronounced tension–compression asymmetry. This is stronger in the medulla, where the compressive peak stress reaches 6.91× the tensile peak compared with 4.12 × in the cortex. Shear is the weakest mode in both regions. The compressive end region steepens sharply after about 5% compression, more so in the medulla than the cortex, with a milder version of the same trend in tension. The medulla data scatter more than the cortex data in absolute terms (Fig. 5). After phosphate-buffered saline immersion, the cortex specimens gain 17.1 % in mean weight against 6.2 % for the medulla (Fig. 6).

**Table 2:**
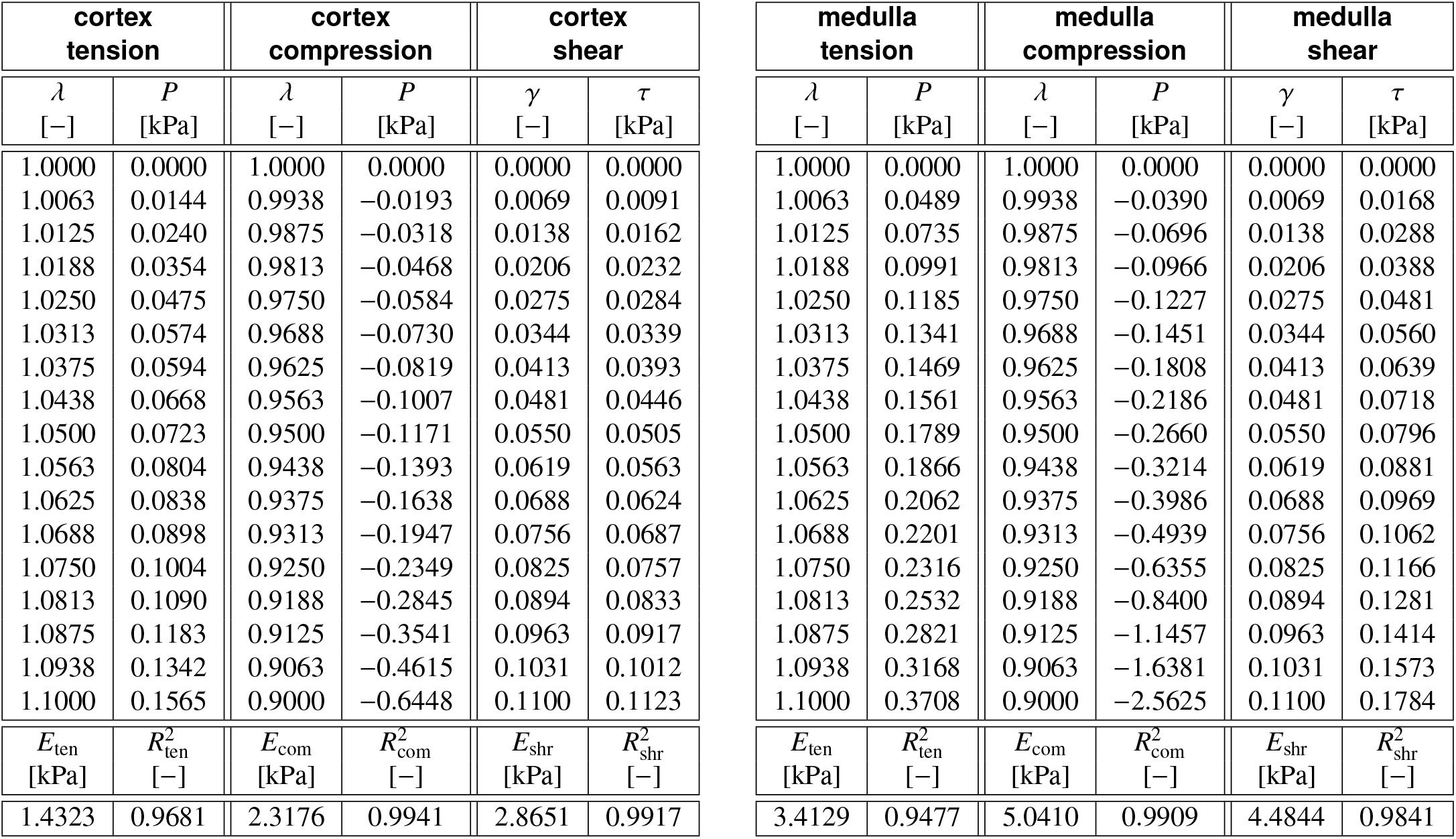
Experimental data and linearized stiffnesses for the cortex and medulla. Stretch–stress pairs for the cortex (left) and the medulla (right) under tension, compression, and shear. Each column gives the specimen mean per mode and region, resampled to 17 equidistant points. Linear moduli *E* are obtained from linear regressions, the coefficients of determination *R*^2^ quantify the goodness of each linear fit.

**Figure 5:**
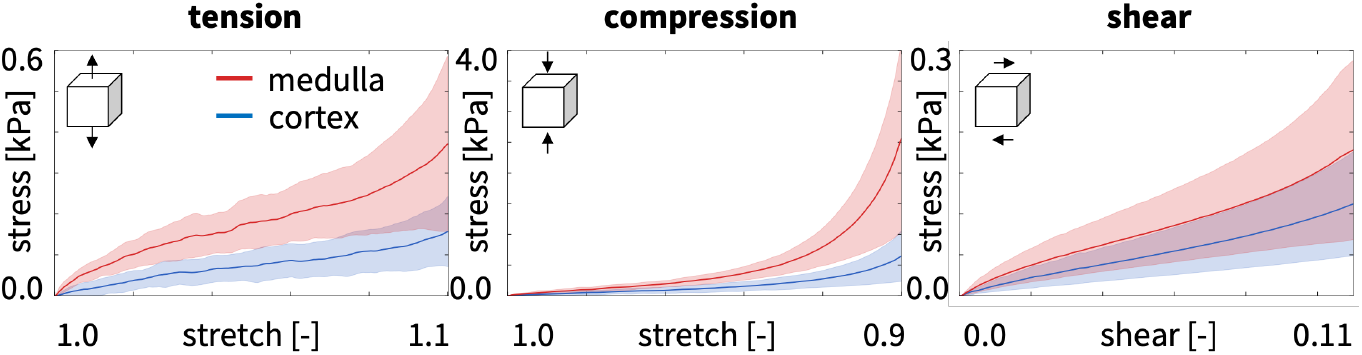
Experimental tension, compression, and shear data for the medulla and cortex. Piola stress versus stretch and shear for the cortex (blue) and the medulla (red) under tension, compression, and simple shear. Solid lines mark the specimen-averaged response and shaded bands the standard deviation. The medulla is stiffer than the cortex across all three modes.

**Figure 6:**
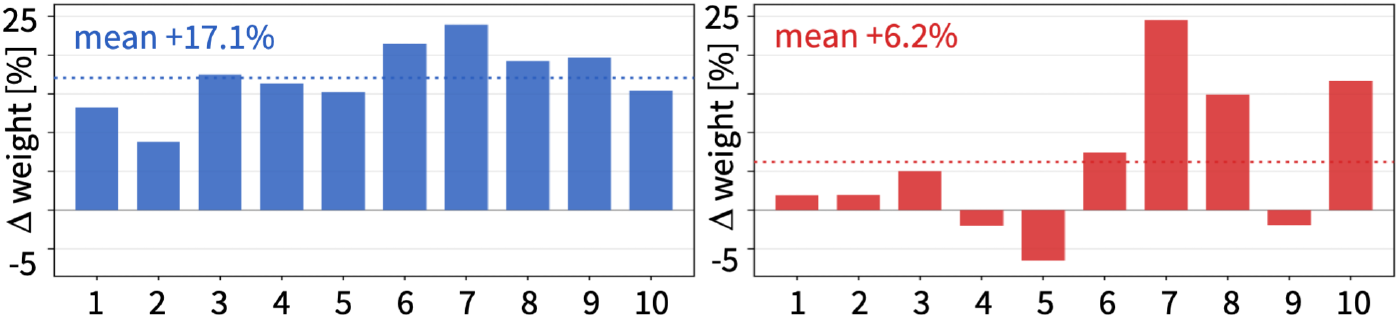
Specimen swelling. Percentage weight change of each specimen after phosphate-buffered saline immersion, with the mean marked by the dashed line. The cortex (blue, left) swells markedly (mean +17.1 %) while the medulla (red, right) swells less (mean +6.2%).

### 3.2. Discovered models

Training on all three modes together using eq. (10), the network discovers the following strain energy functions. For the cortex, the network discovers two terms,

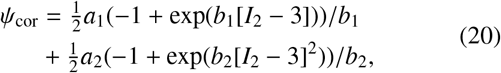

where *a*_1_ ∼ *N* (779 Pa, (556 Pa)^2^), *b*_1_ = 0.18, *a*_2_∼ *N* (111 Pa, (111 Pa)^2^), and *b*_2_ = 4283. For the medulla, the network also discovers an additional linear *I*_2_ term,

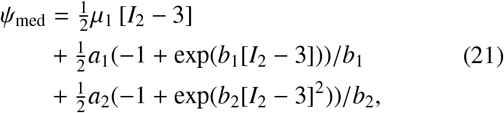

where *μ*_1_∼ *N* (783 Pa, (717 Pa)^2^), *a*_1_ ∼ *N* (822 Pa, (458 Pa)^2^), *b*_1_ = 0.16, *a*_2_∼ *N* (2183 Pa, (2183 Pa)^2^), and *b*_2_ = 3260. Both regions discover a strain energy in *I*_2_ alone, with no *I*_1_ dependence. In both regions the quadratic exponential term carries the smallest weight *a*_2_, yet its contribution rises sharply toward *λ* = 0.9 (Figs. 11 and 13). The quadratic exponential weights *a*_2_ also have standard deviations equal to their means with Σ_*ii*_ → 1. Term selection depends on the training set. The tension-trained, tension+compression-trained, and all-modes models select *I*_2_ terms alone, whereas the compression-trained and shear-trained models discover *I*_1_ terms in both regions (Figs. 11 and 13).

### 3.3. Fit across loading modes and training sets

We evaluate model performance by the excess negative log-likelihood, eNLL (Section 2.6). Trained on all three modes, the model fits the cortex with eNLL = 0.40 in tension, 0.27 in compression, and 0.13 in shear, with a mean of 0.27 across all three modes, and the medulla with eNLL = 0.48 in tension, 0.29 in compression, and 0.12 in shear, with a mean of 0.30 across all three modes (Figs. 7 and 9). For the cortex, cross-mode fit degrades when we train on a single mode and evaluate on the others (Fig. 7, first three columns). Training on compression alone still extrapolates well to shear with eNLL = 0.13, whereas training on tension alone with eNLL= 0.74 does not. Under the all-modes fit (Fig. 7, last column), the model fits shear best with eNLL = 0.13, then compression with eNLL= 0.27, then tension with eNLL= 0.40. The predictive uncertainty tracks the data spread in compression and shear, but exceeds the data spread in tension, and the model mean runs above the data mean across most of the tension range. This pattern is stronger in the medulla (Fig. 9), consistent with its stronger tension–compression asymmetry (Section 3.1). Single-mode training generalizes poorly across the uniaxial modes. Except when we train on compression, the models fail to capture the steep compressive end region. Shear, in contrast, extrapolates well from every training set, with eNLL ≤ 0.17. The model variance tracks the data scatter in the well-fit modes, but in the failed compression extrapolations it stays too narrow to cover the missed mean. Similar to the cortex, the model mean runs above the data mean across most of the tension range.

**Figure 7:**
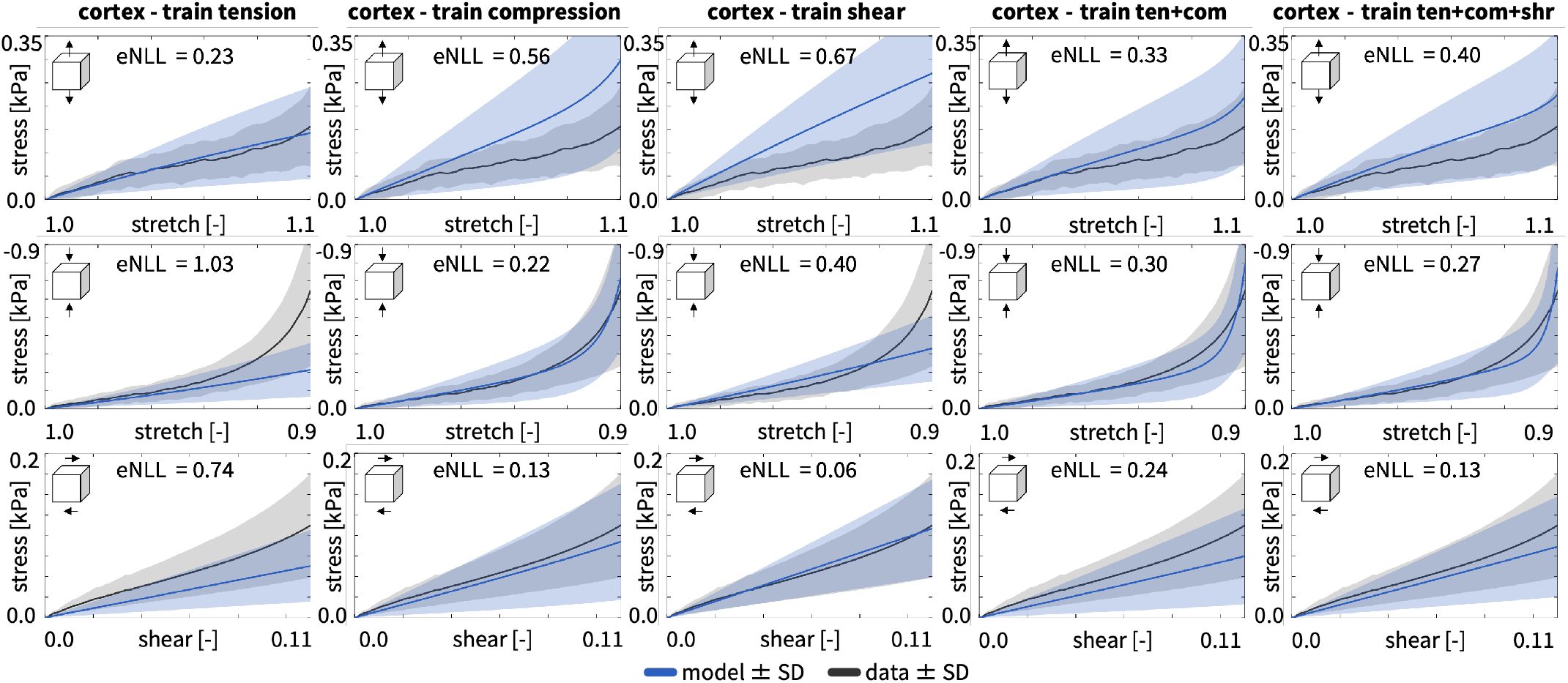
Cortex data and model predictions across modes and training sets. Rows are tension, compression, and shear; columns are the five training sets, tension, compression, shear, tension+compression, and all three modes. Solid lines mark the model mean (blue) and the data mean (gray), and shaded bands their standard deviations. Each panel reports the excess negative log-likelihood, eNLL (Section 2.5).

### 3.4. Enhanced network architecture

The enhanced logarithmic architecture (Fig. 4) trained on all three modes together discovers the following strain energy functions. For the cortex, the network discovers three terms and replaces the linear exponential term of eq. (20) with a linear and a logarithmic term,

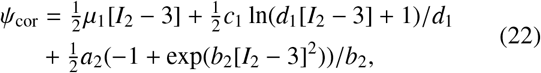

where *μ*_1_ ∼ *N* (394 Pa, (337 Pa)^2^), *c*_1_ ∼ *N* (690 Pa, (690 Pa)^2^), *d*_1_ = 291, *a*_2_ ∼ *N* (3901 Pa, (3398 Pa)^2^), and *b*_2_ = 1229. For the medulla, the network discovers four terms. It keeps the linear exponential and quadratic exponential terms of eq. (21) and replaces the linear term with a logarithmic and a quadratic term,

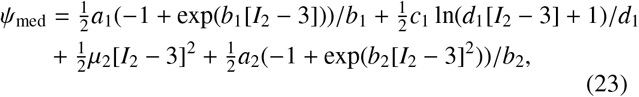

where *a*_1_ ∼ *N* (537 Pa, (327 Pa)^2^), *b*_1_ = 0.71, *c*_1_ ∼ *N* (1673 Pa, (1077 Pa)^2^), *d*_1_ = 355, *μ*_2_ ∼*N* (22378 Pa, (22378 Pa)^2^), *a*_2_ ∼ *N* (2453 Pa, (2453 Pa)^2^), and *b*_2_ = 2823. Both regions again discover a strain energy in *I*_2_ alone, but now also select the logarithmic term (Figs. 12 and 14). The logarithmic architecture fits the joint data better in both regions (Figs. 8 and 10). Trained on all three modes, it has eNLL = 0.26 in tension, 0.11 in compression, and 0.14 in shear for the cortex, with a mean of 0.17 against 0.27, and eNLL = 0.27 in tension, 0.14 in compression, and 0.08 in shear for the medulla, with a mean of 0.16 against 0.30. Tension remains the worst-fit mode in both regions. The improvement does not extend to shear extrapolation. Whenever the training set excludes shear, the enhanced network architecture predicts shear worse than the initial architecture, in both regions and in all three such columns. For the cortex, eNLL rises from 0.74 to 1.13 under tension training, from 0.13 to 0.27 under compression training, and from 0.24 to 0.43 under tension+compression training (Figs. 7 and 8). The medulla follows the same pattern (Figs. 9 and 10).

**Figure 8:**
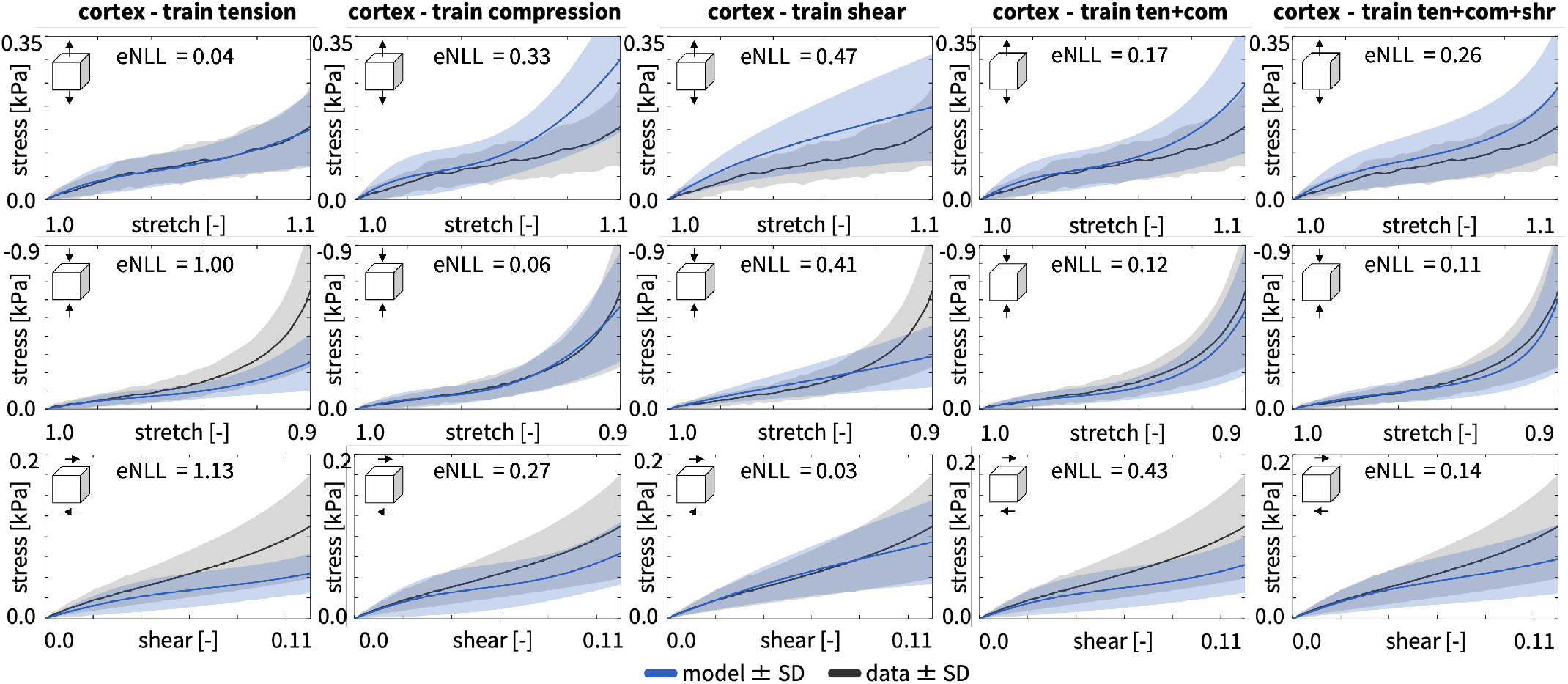
Cortex data and model predictions for the enhanced network. The graphs reports the same experimental data as Figure 7, but the model uses the enhanced network in Figure 4. Rows are tension, compression, and shear; columns are the five training sets, tension, compression, shear, tension+compression, and all three modes. Solid lines mark the model mean (blue) and the data mean (gray), and shaded bands their standard deviations. Each panel reports the excess negative log-likelihood, eNLL (Section 2.5).

**Figure 9:**
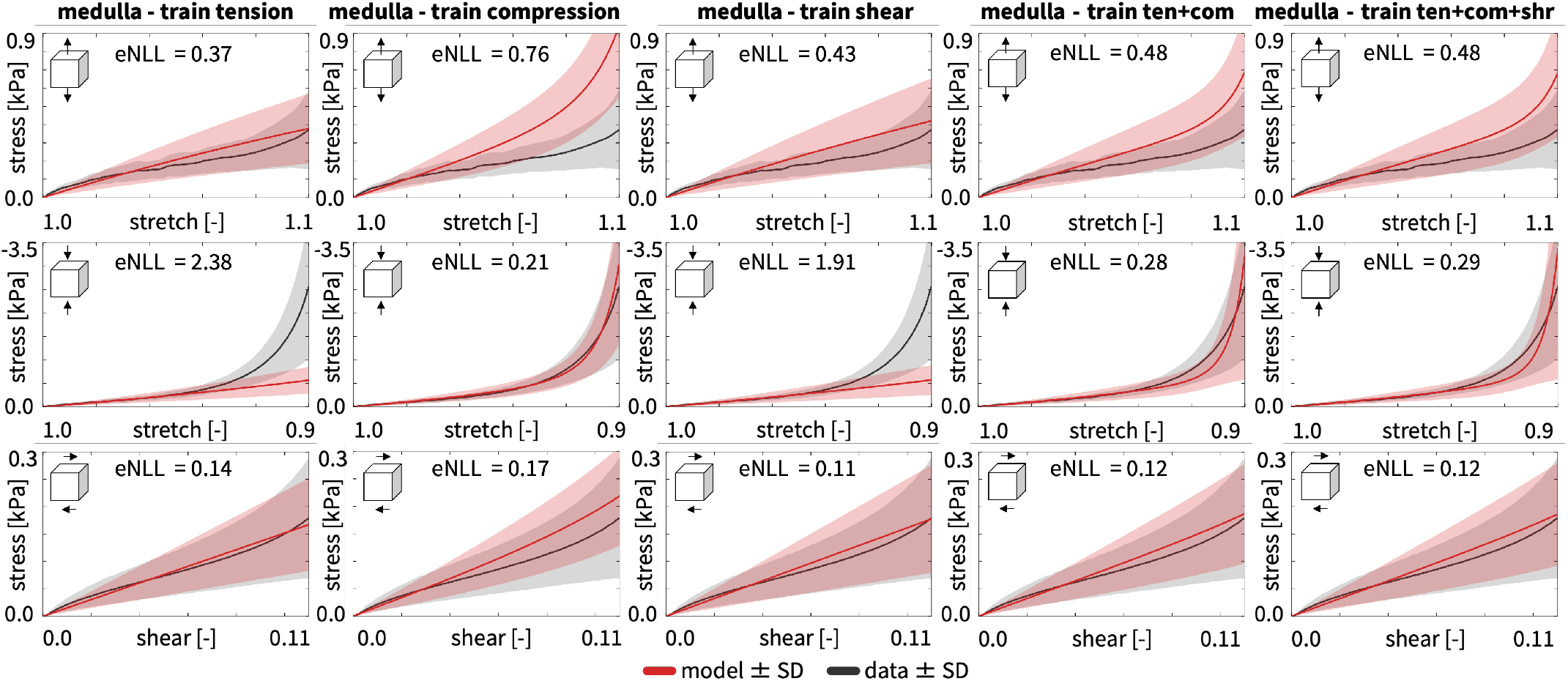
Medulla data and model predictions across modes and training sets. Rows are tension, compression, and shear; columns are the five training sets, tension, compression, shear, tension+compression, and all three modes. Solid lines mark the model mean (blue) and the data mean (gray), and shaded bands their standard deviations. Each panel reports the excess negative log-likelihood, eNLL (Section 2.5).

**Figure 10:**
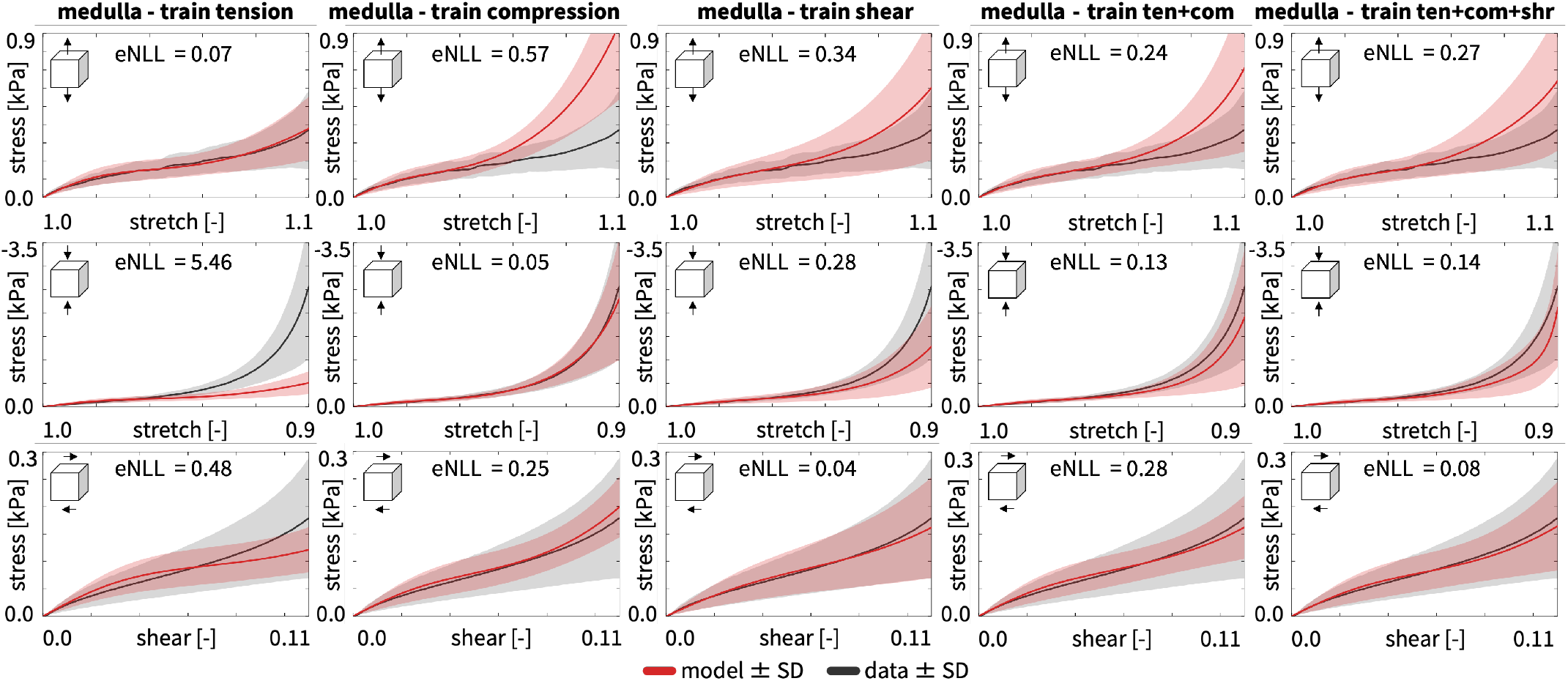
Medulla data and model predictions for the enhanced network. The graphs reports the same experimental data as Figure 7, but the model uses the enhanced network in Figure 4. Rows are tension, compression, and shear; columns are the five training sets, tension, compression, shear, tension+compression, and all three modes. Solid lines mark the model mean (blue) and the data mean (gray), and shaded bands their standard deviations. Each panel reports the excess negative log-likelihood, eNLL (Section 2.5).

**Figure 11:**
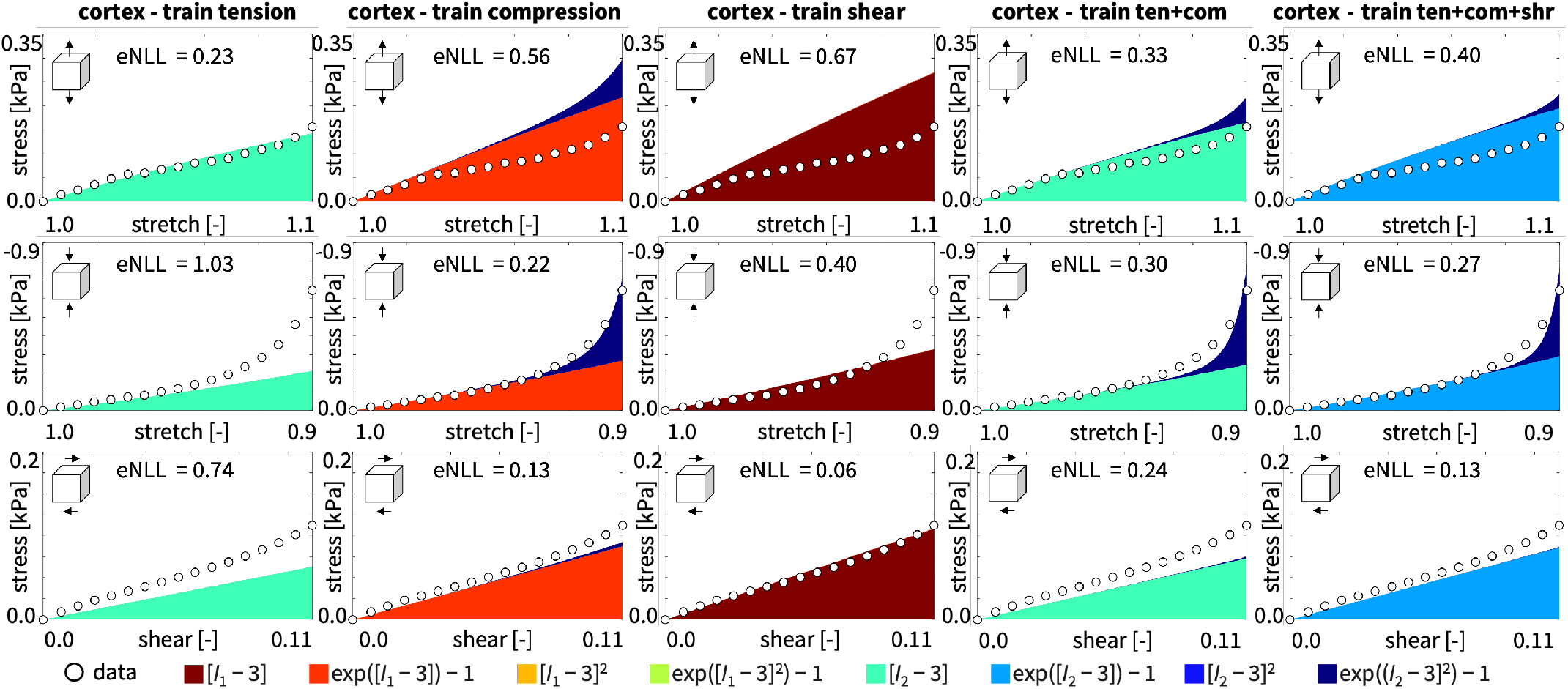
Cortex data and term-wise representation of the discovered strain energy function. White dots show the experimental data mean. Stacked color bands show each discovered term’s contribution to the predicted stress using eq. (10). Rows are tension, compression, and shear; columns are the five training sets.

**Figure 12:**
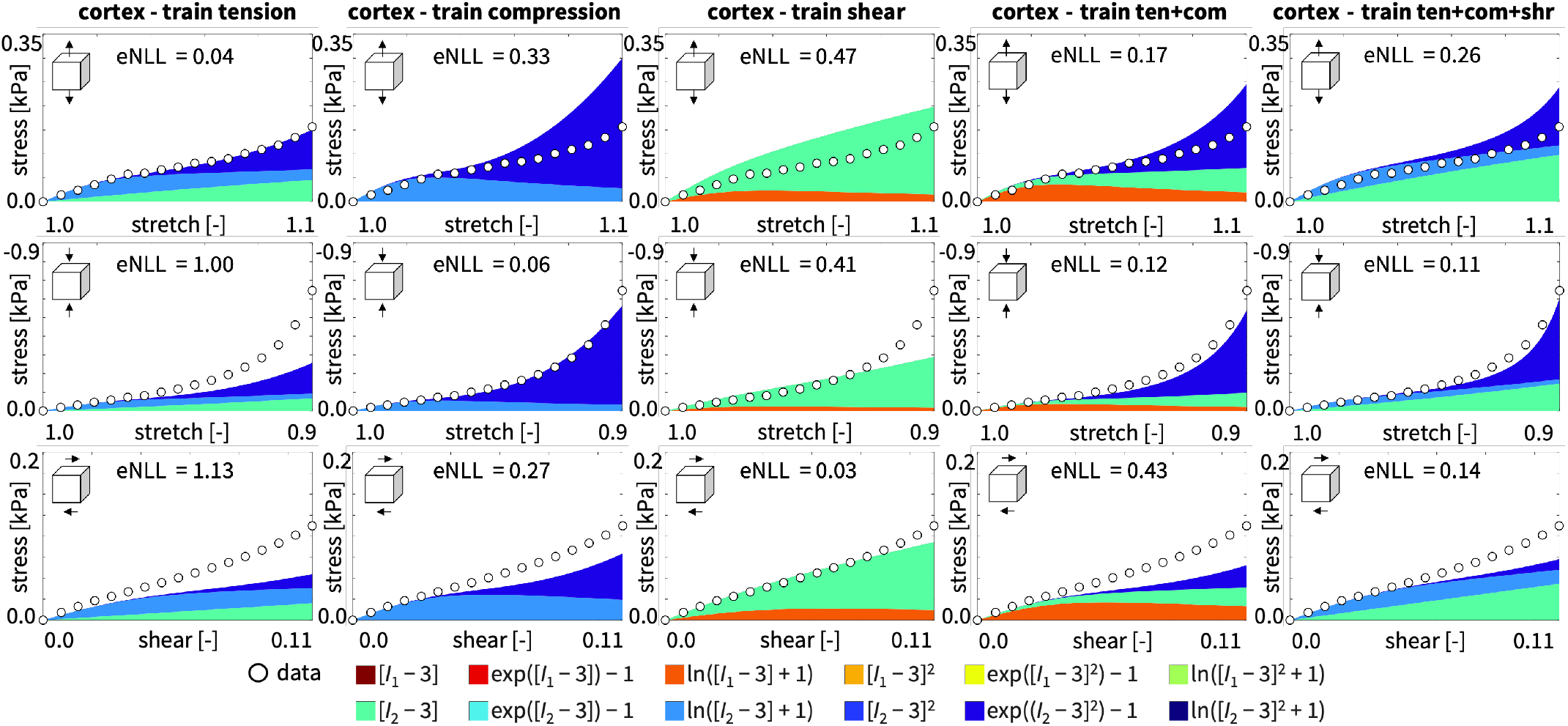
Cortex data and term-wise representation of the discovered strain energy function for the enhanced network. White dots show the experimental data mean. Stacked color bands show each discovered term’s contribution to the predicted stress using eq. (11). Rows are tension, compression, and shear; columns are the five training sets.

**Figure 13:**
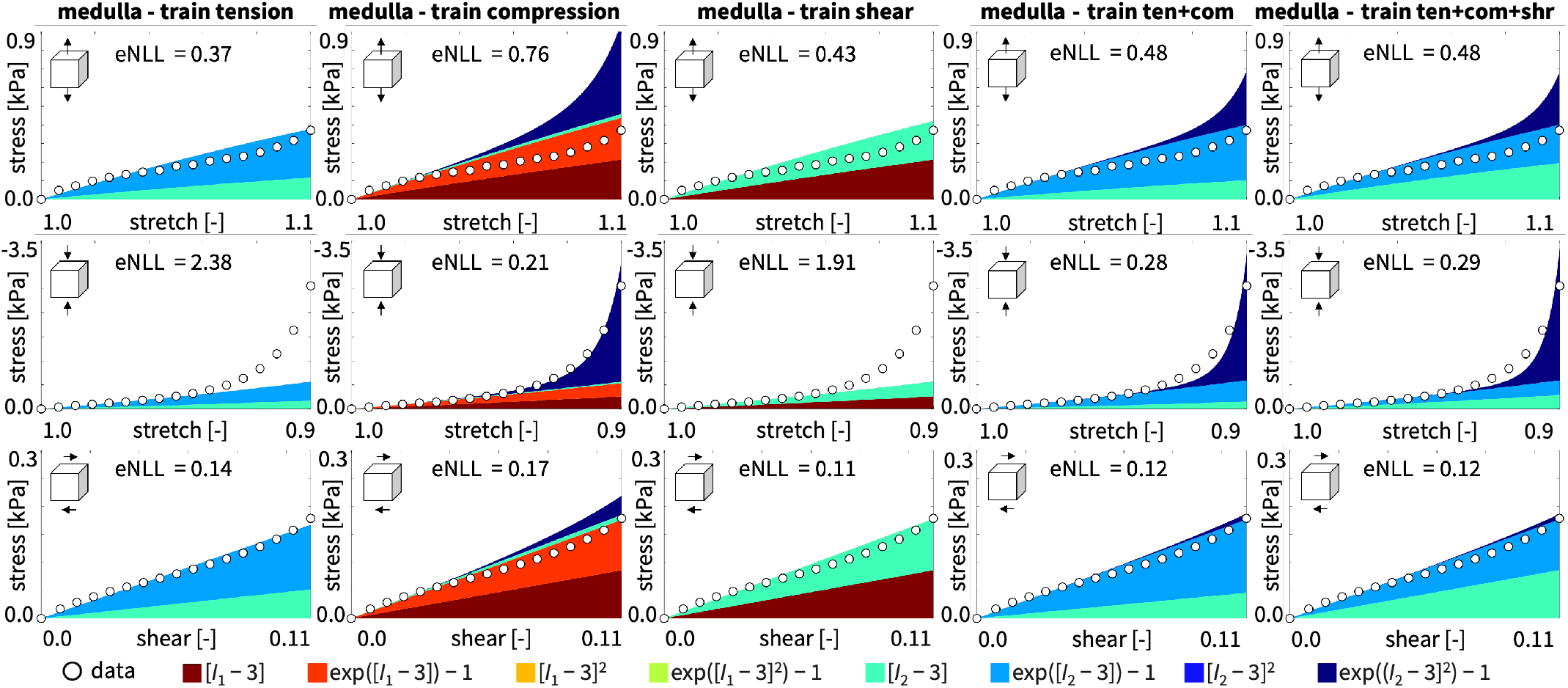
Medulla data and term-wise representation of the discovered strain energy function. White dots show the experimental data mean. Stacked color bands show each discovered term’s contribution to the predicted stress using eq. (10). Rows are tension, compression, and shear; columns are the five training sets.

**Figure 14:**
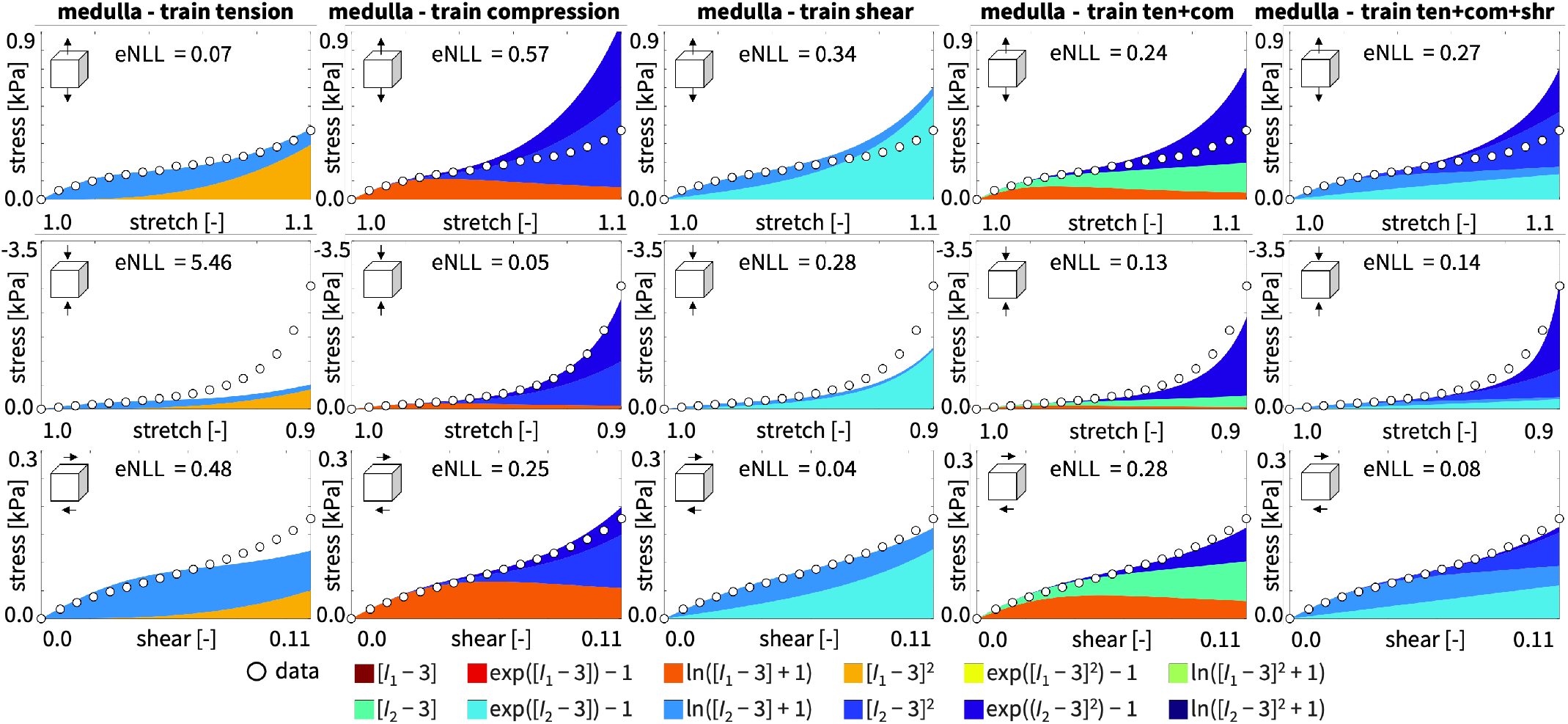
Medulla data and term-wise representation of the discovered strain energy function for the enhanced network. White dots show the experimental data mean. Stacked color bands show each discovered term’s contribution to the predicted stress using eq. (11). Rows are tension, compression, and shear; columns are the five training sets.

## 4. Discussion

We perform tension, compression, and simple shear tests on porcine kidney cortex and medulla. We use a Gaussian constitutive artificial neural network to discover the model, parameters, and uncertainty that best explain these tests. This results in the first kidney model for either region that predicts the stress variance alongside the mean. We report the two-activation architecture (Fig. 3) as the primary model and use the enhanced logarithmic architecture (Fig. 4) as a comparison. The medulla is stiffer in every mode, and the networks discover strain energy functions in *I*_2_ alone for both regions when we train on all three modes.

### 4.1. Regional stiffness of cortex and medulla

Our linearized mechanical response reveals a consistent stiffness contrast between the cortex and medulla. The effective Young’s moduli of the cortex are *E*_ten_ = 1.43 kPa, *E*_com_ = 2.32 kPa, and *E*_shr_ = 2.87 kPa, compared with 3.41 kPa, 5.04 kPa, and 4.48 kPa for the medulla, such that the medulla is 2.38, 2.18, and 1.57 times stiffer in tension, compression, and shear. These values are consistent with the low-kPa range reported for porcine kidney tissue: Shear measurements of porcine renal cortex have reported an elasticity of 1.81 ± 0.17 kPa [2] and a storage modulus of approximately 2.4 kPa at 4 Hz [39], comparable in magnitude to our shear moduli of 0.96 kPa for the cortex and 1.49 kPa for the medulla. Regional shear-wave elastography has further shown that the medulla is stiffer than the cortex [47], consistent with the regional stiffness contrast observed here. Yet, tissue preservation may contribute to differences in absolute stiffness values [43]. Taken together, literature supports the range of our linearized moduli, while our direct mechanical tests reveal a consistent regional contrast across all three loading modes, with the medulla approximately twice as stiff as the cortex. This pronounced mechanical heterogeneity can strongly influence how deformation and stress redistribute between cortex and medulla during surgical manipulation, needle interventions, and renal trauma.

### 4.2. Discovered models

When trained on all three modes, for both regions, the network discovers strain energy functions in *I*_2_ alone, with no *I*_1_ dependence. To date, constitutive modeling of kidney tissue has been based on selecting a fixed functional form and fitting its parameters to data. As mentioned in Section 1, past hyperelastic forms include Blatz-Ko [11] and Mooney–Rivlin [36, 49]. Here the data themselves select it. Interestingly, human brain tissue shows the same preference for the second invariant *I*_2_ [27], and mechanistic considerations link this to a shear-dominatant behavior—the Poynting effect—and a stronger tension–compression asymmetry than *I*_1_ can produce [23]. Our data display the same asymmetry (Section 3.1), and the network expresses a preference for *I*_2_.

### 4.3. Identifiability

Several researchers caution against *I*_2_-only models and attribute their discovery to limited experimental data [4]. Over the tested range, [*I*_1_ −3] and [*I*_2_ −3] differ at most by ten percent and the linear and linear-exponential terms are almost indistinguishable. Selecting *I*_2_ over *I*_1_ also depends on the training set. The compression-trained and shear-trained models discover *I*_1_ terms in both regions (Section 3.2, Figs. 11 and 13). In shear *I*_1_ = *I*_2_, so the choice is arbitrary. We conclude that the *I*_2_-only model describes the data well over the tested range, but how it performs at larger stretches remains to be studied in more detail.

### 4.4. Generalization across loading modes

The models discovered for tension or shear only do not generalize well to predict the compressive behavior. The steep compressive end region near *λ* = 0.9 illustrates this misfit most clearly (Section 3.3). The cortex model generalizes to compression better than the medulla, whose stronger tension–compression asymmetry is a possible cause. Shear behaves in the opposite way. The models predict it well without training on it, and adding it to the training set changes the uniaxial predictions little.

### 4.5. Predicted uncertainty

The predicted uncertainty band exceeds the measured scatter in tension in both regions and misses the mean in the failed compression extrapolations (Section 3.3). Two features of the formulation limit how far the band can follow mode-specific scatter. First, the predicted variance inherits the term structure of the mean, since eq. (15) scales each term by its own relative spread *w*_*σ,i*_. The same term dominates tension and shear in both regions, so a spread wide enough to cover the shear scatter widens the tension band as well. Second, eq. (18) divides the squared residual by the variance, so a mode whose mean the network fits poorly rewards a wider band. Tension is the worst-fit mode in both regions and the network overestimates its mean (Section 3.3), so both mechanisms push the increase in tension variance. In both cortex and medulla, the quadratic exponential weight *a*_2_ has a standard deviation equal to its mean, so *Σ*_*ii*_ sits at the constraint boundary of eq. (16). This observation agrees with previous studies that attribute it to a stress variance that is large relative to the mean [31]. Our compressive data suggest a second mechanism. The measured scatter stays small over most of the stretch range and grows sharply toward *λ* = 0.9, where the quadratic exponential term governs the response (Section 3.2). Two interpretations follow. In the first, the term absorbs genuine scatter, the constraint binds, and the reported standard deviation understates the spread. In the second, the term contributes so little to the predicted variance over most of the range that the likelihood barely constrains *w*_*σ,i*_, and its value drifts to the boundary. Since these two interpretations imply opposite corrections, we treat every weight at the boundary as weakly identified.

### 4.6. Architecture comparison

In both regions, the enhanced network (Fig.4)) improves the joint fit compared to the simple network (Fig. 3) when we train on all three modes. The gain lies in the uniaxial modes, while shear changes little (Section 3.4). The logarithmic term adds a shape compared with the other activations. This reproduces the initial toe of the tension and compression data that the convex terms miss. Yet, whenever the training set excludes shear, the enhanced predicts shear worse than the simple network. When trained on all three modes, both architectures discover strain energy functions in *I*_2_ alone for both regions. The additional logarithmic terms change the quality of the fit, but not the invariant class of the discovered model. The logarithmic terms are concave, so eq. (11) no longer guarantees polyconvexity, and concave terms can in principle produce decreasing stress. Here, the discovered models increase monotonically across the entire tested regime in every mode. Any use of free energy functions with logarithmic terms eq. (11) should confirm monotonicity within the relevant range. We therefore report the linear/linear-exponential architecture as the primary model, since it always guarantees polyconvexity in the mean (Fig. 3).

### 4.7. Limitations

While our discovered models show promising results, our approach has a few limitations that point towards future studies: First, we model both regions as isotropic. The assumption is reasonable for the cortex, but the directional microstructure of the medulla may violate it [47]. Second, we purchase the kidneys from a commercial grocer, so the exact post-mortem time before testing is unknown. Since the post-mortem time and freeze–thaw may alter the mechanical response [43], future studies should use fresh kidneys with a recorded post-mortem time. Third, testing at room temperature, gluing specimens to the rheometer, and using a fixed compression–tension–shear sequence may have affected the mechanical response. Fourth, we define the elastic response as the average of the loading and unloading branches. Our quasi-static hyperelastic treatment does not represent the hysteresis we record between cycles, and future studies should consider modeling the kidney as hyperviscoelastic [16]. Fifth, we treat the kidney as a single-phase hyperelastic solid and neglect fluid–solid interaction and fluid motion, which we could address by switching to a poroelastic or poroviscoelastic model [5].

## 5. Conclusion

Existing constitutive models of the kidney prescribe the strain energy function a priori, treat tissue variability deterministically, and do not distinguish between cortex and medulla. Here we perform tension, compression, and shear tests on the cortex and medulla, and train a Gaussian constitutive artificial neural network to discover the strain energy function, its parameters, and its predictive uncertainty for both regions. We observe that the medulla is approximately twice as stiff as the cortex across all three loading modes. The effective Young’s moduli are 3.41, 5.04, and 4.48 kPa for the medulla and 1.43, 2.32, and 2.87 kPa for the cortex in tension, compression, and shear, corresponding to stiffness ratios of 2.38, 2.18, and 1.57. Both regions display pronounced tension–compression asymmetry, which is stronger in the medulla. When trained on all three modes combined, our network discovers strain energy functions in the second invariant alone, with no dependence on the first invariant over our tested deformation range. The cortex model features two exponential *I*_2_ terms, while the medulla model includes an additional linear *I*_2_ term. Our Gaussian external weights propagate specimen-to-specimen variability into closed-form probabilistic stress predictions. Adding logarithmic terms to the network improves the fit across all three modes, but reduces the ability to predict unseen shear tests. Since these logarithmic terms no longer guarantee polyconvexity, we recommend the simple linear-exponential network for model discovery. Our discovered probabilistic models of the cortex and the medulla provide region-specific constitutive laws that capture the mechanical heterogeneity of the kidney and propagate tissue variability into uncertainty bounds. These advances enable more realistic predictions of kidney deformation and stress for surgical planning, needle interventions, and renal trauma, while establishing a healthy-tissue baseline for the mechanical quantification of renal disease.

## CRediT authorship contribution statement

VE: Conceptualization, Methodology, Software, Validation, Formal analysis, Investigation, Data curation, Writing–original draft, Writing–review & editing, Visualization; SRStP: Conceptualization, Methodology, Formal analysis, Writing–review & editing; EK: Conceptualization, Methodology, Formal analysis, Writing–review & editing.

## Data availability

Our source code, data, and examples are available at https://github.com/LivingMatterLab/CANN.

## Acknowledgments

This research was supported by the Aker Scholarship to VE, by the NSF Graduate Research Fellowship and the Stanford Diversifying Academia, Recruiting Excellence Fellowship to SRStP, and by the NSF CMMI Award 2320933 and the ERC Advanced Grant 101141626 to EK.

